# Shrinkage of dispersion parameters in the binomial family, with application to differential exon skipping

**DOI:** 10.1101/012823

**Authors:** Sean Ruddy, Marla Johnson, Elizabeth Purdom

## Abstract

The prevalence of sequencing experiments in genomics has led to an increased use of methods for count data in analyzing high-throughput genomic data to perform analyses. The importance of shrinkage methods in improving the performance of statistical methods remains. A common example is that of gene expression data, where the counts per gene are often modeled as some form of an over-dispersed Poisson. In this case, shrinkage estimates of the per-gene dispersion parameter have led to improved estimation of dispersion in the case of a small number of samples.

We address a different count setting introduced by the use of sequencing data: comparing differential proportional usage via an over-dispersed binomial model. This is motivated by our interest in testing for differential exon skipping in mRNA-Seq experiments. We introduce a novel method that is developed by modeling the dispersion based on the double binomial distribution proposed by Efron (1986). Our method (WEB-Seq) is an empirical bayes strategy for producing a shrunken estimate of dispersion and effectively detects differential proportional usage, and has close ties to the weighted-likelihood strategy of edgeR developed for gene expression data (Robinson and Smyth, 2007; Robinson *et al.*, 2010). We analyze its behavior on simulated data sets as well as real data and show that our method is fast, powerful and gives accurate control of the FDR compared to alternative approaches. We provide implementation of our methods in the R package `DoubleExpSeq` available on CRAN.

## 1 Introduction

In genomic studies, a common approach to high dimensional data is to marginally examine the effect of each feature with a simple statistical test in order to find the most promising features. A well-known example of this type of marginal testing is that of gene expression studies, where the features of each sample consists of measurements of the mRNA levels of tens of thousands of genes from the sample. In this setting, there are generally few samples (on the order of 10 or less) and thousands of features or genes. A common analysis is to perform a statistical test separately for each gene; for example, a t-test per gene to determine if there is a difference between two groups of samples. In such a paradigm, it has been found that shrinkage of the individual parameter estimates or test statistics greatly improves the results, and a great deal of work has been done in different settings to this purpose.

The setting for many of these shrinkage routines was initially in the context of continuous, roughly log-normal intensity data from microarray experiments. The growth of relatively cheap sequencing technologies has resulted in sequencing becoming preferred over the previous generation of microarray technologies. The result of sequencing experiments, unlike the continuous intensity measurements of microarray experiments, is generally a count of the number of sequences matching a criterion, such as the number of sequences from a particular gene. As a result there has been great interest in how to most effectively use discrete distributions for common tasks that previously relied on normal data. For the marginal testing approaches, this includes appropriate use of over-dispersed models and how to provide shrinkage methods.

A common type of question in sequencing experiments is to compare the counts of sequences measured across different conditions, such as the two group setting described above. One setting in which this is frequent is the case of finding differences in the mRNA levels of a gene in different conditions. In this case, the counts per gene are the number of sequenced mRNA that come from that gene. There are different amounts of total sequences collected from different samples, so that the question of interest is whether the proportion of counts allocated to a given gene varies across conditions. In the gene setting, however, the total number of sequences in a sample is in the millions and are spread across thousands of genes, so that the proportions are quite small. For this reason it is common to use a Poisson distribution to model the counts with an offset parameter equal to the total number of sequences (Marioni *et al.*, 2008). Generally an over-dispersed model is preferred, and the prominent modeling technique for over-dispersion has been to use a negative-binomial (Robinson and Smyth, 2007), though some methods have incorporated over-dispersed binomial distributions such as the beta-binomial. For all of these gene expression methods, there has also been focus on creating shrinkage estimators of the dispersion parameter which greatly improve the performance of the methods in small sample sizes (Robinson and Smyth, 2007; Anders and Huber, 2010; Zhou *et al.*, 2011; Yang *et al.*, 2012; Wu *et al.*, 2013; Yu *et al.*, 2013; Leng *et al.*, 2013).

We are interested in a slightly different setting, namely when we are performing marginal testing comparing proportions but the Poisson approximation is not valid and the proportions can take on the full range of 0 to 1. We are motivated by the the question of detecting differences in alternative splicing between conditions – specifically an approach that simplifies the problem by focusing on each exon separately and evaluating whether it is excluded (“spliced out”) more frequently in some conditions than others. We focus on the measure of exclusion often called *proportion spliced in* (PSI or ψ) that measures the number of sequences that show inclusion of the exon compared to the number excluding or skipping the exon (Pan *et al.*, 2008; Venables *et al.*, 2009; Wu *et al.*, 2011; Shen *et al.*, 2012; Barbosa-Morais *et al.*, 2012; Brooks *et al.*, 2014). There are alternative approaches to summarizing mRNA-Seq data for detecting differential alternative splicing, but focusing on PSI per exon has the advantage that it is a relatively simple summary of the data that also makes use of the large amount of information available in junction reads that span introns (also called split-reads). We describe and give the necessary background to this problem in Section 2 below.

Our methodological focus, like that of the more standard gene expression setting, is on shrinkage estimates of the dispersion parameter, which is a common focus of high-throughput genomic settings. Unlike the gene expression example, PSI values can take the full range of values from zero to one, and therefore existing work based on over-dispersed Poisson models from gene expression are not applicable in this setting.

We propose a novel empirical bayes framework for estimating the dispersion parameter for data whose distribution is part of the family of dispersed exponential distributions proposed by Efron (1986), called the double exponential family of distributions. Our empirical bayes method framework is based on the fact that the distribution of the dispersion parameter can be shown to be approximately Gamma distributed, which we develop below. In addition to being a tractable distribution, the estimates produced from the double exponential family model have close ties to quasi-likelihood estimates which are widely used for estimation of binomial over-dispersion. Given this close connection, our method is effectively an empirical bayes method for quasi-likelihood estimation of the dispersion parameter.

Our empirical bayes framework provides two, related versions. The first is the standard empirical bayes estimator (DEB-Seq). The second (WEB-Seq) is also an empirical bayes estimator with a different parameterization of the prior; it is related to the weighted likelihood method of shrinkage of Robinson and Smyth (2007) applied to the double exponential family of distributions, but unlike that approach, the empirical bayes methodology provides a data-driven estimate for the tuning parameter.

We compare the performance of our method to other methods and demonstrate that in addition to providing a fully automated method for shrinkage, our methods have superior performance on simulated data in the exon inclusion setting. We also apply these methods to mRNA-Seq data from real tumor samples generated by the Cancer Genome Atlas project (Cancer Genome Atlas Research Network, 2011) which suggests that it can similarly control the false discovery rate and find promising targets of splicing. Furthermore, there is very little computational overhead in our methods compared to many existing methods.

## 2 **Differential Exon Usage**

### 2.1 **Alternative Splicing**

The static genetic code found in the DNA of each cell must be read and converted into the molecular products that are used throughout the cell, e.g. proteins. Specifically, a portion of the DNA, called a gene, encodes which specific amino acids will make up a protein. When a protein is needed, regulatory processes in the cell trigger its corresponding DNA to be transcribed into an independent copy of the DNA, called mRNA; the genetic code contained in the mRNA is then translated into the string of amino acids that form the protein.

In eukaryotic cells, the process of copying the DNA into mRNA itself has stages. A direct copy of the DNA is created, called a pre-mRNA, and the pre-mRNA is then further processed into an mRNA. This always involves some changes to the pre-mRNA (e.g. adding a poly-A tail and a 5’ cap), but in many eukaryotes, the processing of pre-mRNA goes further and selectively cuts out portions of the pre-mRNA so that the code for the protein contained in the final (mature) mRNA transcript is not an exact copy of the code found in the DNA of the cell. The process of cutting out portions of the mRNA product is called *splicing*.

Splicing is not haphazard but a highly regulated process: the DNA code itself indicates the portions of the pre-mRNA product that should be spliced out and recruits the protein complexes that actually do the splicing. In many complex organisms, including human and many common model organisms like fruitflies and mice, the splicing of a pre-mRNA product is more complex because which parts of the pre-mRNA are removed can vary based on the environmental cues of the cell. The end result is that a single gene representing a stretch of code on the DNA can result in a diversity of mRNA transcripts. This is referred to as *alternative splicing* (again, see Figure 1a for an illustration)

**Figure 1:**
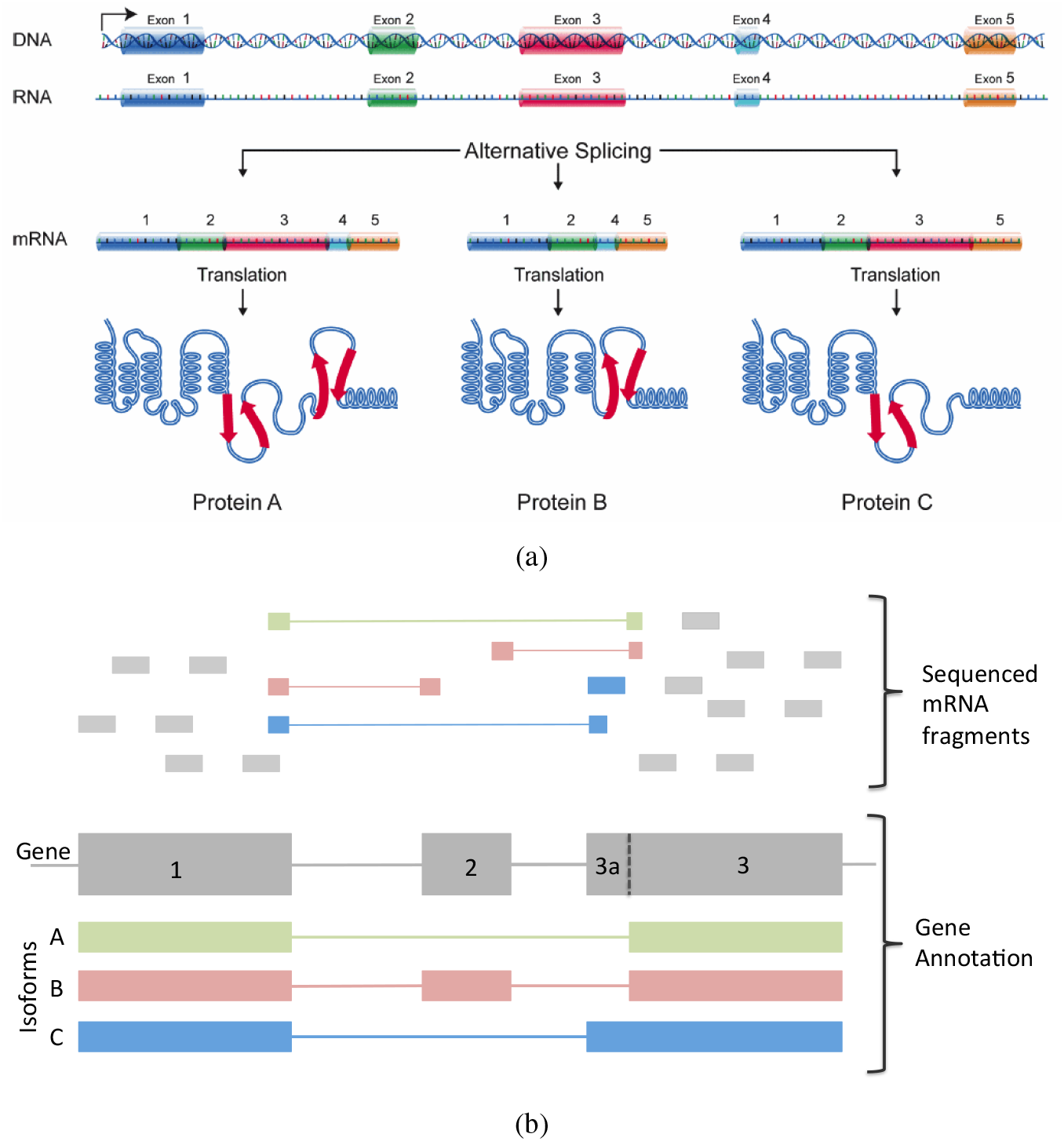
Illustration of Alternative Splicing and Sequencing. (a) An illustration of the process of creating multiple isoforms from a single gene (DNA). A RNA copy of a gene is created (pre-mRNA) that contains all the gene’s exons and introns. The pre-mRNA can then be spliced in one of three ways by removing different combinations of exons, and each possibility leads to a different mRNA (isoform) and protein product. Illustration courtesy of National Human Genome Research Institute (2014) (b) A pictorial example of aligning sequenced fragments when there are multiple isoforms. The bottom boxes represent the traditional annotation of a gene region: the dark grey exons represent all known exons (i.e. the gene), the colorful boxes represent the three isoforms from the gene which have different combinations of the exons. We have divided the exons of the gene into 4 regions, corresponding to the isoforms in which they are contained. The top set of small boxes represents the sequenced fragments. Most reads are represented as small boxes, meaning they completely match the part of genome they are above. Some reads are split into smaller boxes with connecting lines; these represent *junction reads* where the two parts of the fragment match disconnected regions of the genome. The sequenced fragments are also colored by the isoform to which they can be uniquely identified, with the majority of fragments being grey because it is not possible to identify the isoform from which they originate.

The different possible transcripts that can result from a single gene are often called *isoforms* of the gene. The DNA of a gene can be thought of as being divided into *introns* (the portion that will be removed) and *exons* (the portions that will remain), though because of alternative splicing some regions may be retained in one isoform and removed in another isoform. How many isoforms there are in a gene and how interrelated the isoforms are varies by gene. Determining the possible isoforms based on the DNA of a gene is difficult, and the set of all possible isoforms is not completely determined even for well characterized genomes like human.

### 2.2 **Sequencing of mRNA for Alternative Splicing**

Previously, large-scale quantification of the amount of mRNA in a cell (also called *mRNA expression*) was via microarray technology. Microarrays allowed an unprecedented ability to measure the mRNA from thousands of genes simultaneously. However, gene expression measurements from microarrays quantified all mRNA from a gene without distinguishing between different isoforms of the gene.

“Next-generation” sequencing technologies have resulted in a rapid decline in the price of sequencing and allow for widespread sequencing of mRNA for expression studies instead of microarrays. Unlike microarrays, the sequencing of mRNA can detect mRNA in a cell without any prior information as to what isoforms exist. Due to this, the sequencing of mRNA provides a great deal of information about alternative splicing in the cell.

While current sequencing methods allow for direct sequencing of mRNA, they still do not allow for an entire mRNA transcript to be sequenced. Instead the mRNA must be cut into small fragments before sequencing, meaning that the sequences that are obtained are only a portion of the mRNA from which they came. If the mRNA has undergone splicing, then the mRNA fragment may not directly match the genome. Usually it is possible to match a fragment back to the genome, where gaps in the match of the mRNA to the genome will correspond to introns that were removed due to splicing (see for example Trapnell *et al.* (2009), Wu and Nacu (2010)). Figure 1b gives an illustration.

The data from mRNA-Seq consisting of these aligned reads can be simplistically thought of as a mix of two different types of sequenced fragments. One set consists of fragments that span two or more exons that were spliced together during the process of removing the introns. As a result they do not directly map to the genome, but rather portions map to one exon and portions map to another; such reads are often called *junction reads* or split-reads (Figure 1b shows examples). Junctions reads are often interesting for alternative splicing because they can give direct evidence of splicing. For example, in Figure 1b, isoforms A and B differ because one includes an exon and the other does not (called an exon skipping event); then fragments that connect exon 1 to exon 3 without including the middle exon (exon 2) must come from isoform A, while those fragments that connect exon 2 to the adjacent exons 1 and 3 must come from isoform B.

Other fragments are contained entirely within an exon. These are less likely to provide direct evidence as to the isoforms from which they came (though we give an example in Figure 1b where there is such a read contained entirely within an exon that must come from isoform C). However, the quantity of reads mapping to different regions can provide indirect evidence of alternative splicing. For example, in the case of the skipping of exon 2, described above, a reduction in isoform B compared to isoform A would result in less total expression of exon 2 (i.e. less reads mapping to it) relative to exons 1 and 3. Since the bases just around the junctions between exons represent only a small portion of mRNA, junction reads are much less common than reads mapping to the exons, and therefore exon reads are important for accurate quantification the the overall expression of the mRNA.

### 2.3 **Measuring Differential Alternative Splicing**

There are different ways to summarize the information in the sequenced fragments with respect to detecting differences in alternative splicing. They can be roughly thought of as making different use of fragments mapping to the junctions versus the interior of the exons. Depending on how the information is summarized, different statistical approaches are appropriate. Our methods here focus on analyzing data resulting from one particular approach to summarizing the information regarding alternative splicing. This method makes use of the junction reads or ‘split reads’ and we call this an inclusion-exclusion summarization, with the summary statistic often called the *percent spliced in* or *exon inclusion percentage*.

The idea of the inclusion-exclusion approach is to break the isoforms into individual contrasting patterns of interest and count how many fragments agree with one pattern compared to another. An example is that of the comparison of isoform A to B described above. In this case, junctions that skip exon 2, joining exons 1 and 3, show evidence of isoform A while those that include exon 2 show evidence of isoform B. This ignores any other exons in the gene, only considering the three exons relevant for assessing the exon skipping event. Then the proportion of these fragments including exon 2 results in a proportion or percentage which is called the percent spliced in for exon 2, abbreviated PSI or ψ. More generally, it is common to break each gene into well-defined simple alternative splicing events (e.g. skipping exons, alternative 5’/3’ splice starts) and evaluate whether the PSI changes between different conditions across all genes (see Pan *et al.* (2008); Venables *et al.* (2009); Wu *et al.* (2011); Shen *et al.* (2012); Barbosa-Morais *et al.* (2012); Brooks *et al.* (2014) for examples). Programs like MATS (Shen *et al.*, 2012), JuncBase (Brooks *et al.*, 2011), DiffSplice (Hu *et al.*, 2013), and SpliceTrap (Wu *et al.*, 2011) take an annotation file of the transcriptome and create counts for these types of alternative splicing events.

In our data analysis, we consider PSI values defined more broadly for other types of events than just these classical definitions of splicing. These classical alternative splicing events are difficult to identify computationally for the whole genome, and they also do not include many other more complicated events. For example, in our simple illustration, there is also isoform C that skips exon 2 but also has an alternative start site for exon 3. In our computations with real data, we calculate a PSI for every single exon, without associating an exon with a type of splicing. We define the fragments that include an exon to be all those contained in the exon, and we define those that exclude the exon to be any junctions that skip the exon regardless of what other exons that junction combines together. So that in our simple illustration, all reads that skip exon 2 would be considered “excluding” exon 2, regardless of whether they originated from isoform A or C. To make the comparison across exons of different lengths more comparable, we can also limit the fragments “including” an exon to just those fragments that join the exon to other exons (e.g. only fragments that overlap exon 2 AND another exon). Either way, this creates a measure of PSI for every exon in the annotation, which we find more satisfying than limiting to just exons that are in simple types of combinations.

We emphasize that regardless of how an ‘event’ is defined, all of these inclusion-exclusion summarizations result in the same kind of data structure from a statistical point of view: *Y* successes out of *m* trials identified for each event.

There are other ways to summarize mRNA-Seq data in order to compare alternative splicing between samples. Our methods are not generally appropriate for these types of summaries (except as noted below), but we mention them here so as to make clear the distinction.

**Isoform Estimation** A direct way to summarize the information in a gene is to estimate the individual isoform estimates directly. Because there is not a direct mapping of most reads to isoforms, this is a deconvolution problem, where the observed expression of a fragment is a convolution of the expression of the individual isoforms of which it could be a part (Denoeud *et al.*, 2008; Jiang and Wong, 2009; Trapnell *et al.*, 2010; Richard *et al.*, 2010; Salzman *et al.*, 2010; Katz *et al.*, 2010). In estimating the isoform expression level, all of the reads of the gene are used. Individual isoform expression levels can be individually compared in a similar way to gene expression estimates (see EBSeq (Leng *et al.*, 2013) which provides for corrections specific to isoform analysis), or specific features of the isoforms can be compared for differential isoform usage (e.g. rSeqDiff (Shi and Jiang, 2013)).

Furthermore, some researchers use isoform expression measures to create PSI values per exon, where instead of using inclusion-exclusion *counts*, they use the isoform measurements to calculate, per exon, the percentage of the overall isoform expression that comes from isoforms including the exon (Shi and Jiang, 2013). So in our simple example, exon 2 is contained only in isoform B, and the PSI for exon 2 would be the isoform abundance for isoform B as a fraction of the total (sum) abundance for the gene (this estimates a similar quantity to our preferred way of calculating PSI per exon described above). Since isoform estimates are continuous, and not counts, this form of a PSI would not have the discrete statistical structure of *Y* successes out of *m* trials; however it is common to convert isoform expressions into expected counts, in which case the expected counts would have that discrete nature.

The ability to estimate isoform expression levels depends on having complete knowledge of all possible isoforms in the gene (called the transcriptome), as well as having enough distinguishing information between the isoforms to deconvolve the isoform expression from the overall expression. There are still many organisms of interest to researchers that do not have extensive knowledge of the transcriptome. While there are computational methods that attempt to construct the set of isoforms de novo, based on mRNA-Seq reads (Trapnell *et al.*, 2010; Guttman *et al.*, 2010; Richard *et al.*, 2010), this is an extremely complicated problem, and these de-novo methods can be unreliable and unstable if used on a single, small experiment or without sufficient numbers of sequenced fragments. For this reason, it can be preferable to use alternative methods of summarizing the data for detecting differential alternative splicing.

**Relative Exon Usage** Another approach to alternative splicing is to evaluate the relative expression of exons to detect alternative splicing, ignoring the specific information in the junction reads. As we noted above, if isoform B is less expressed, this should be apparent in relatively less total number of counts overlapping exon 2; in theory this can be done without any recourse to what specific exons are joined together by junction reads. For relative exon usage, the input per exon is all counts overlapping an exon. Relative exon usage does not make use of the information of how junction fragments skip the exons, except in their contribution to reads overlapping an exon.

In practice, many technical aspects of sequencing can create biases so that some exons are more likely to get sequenced reads affecting the overall count of an exon relative to other exons in the gene. For this reason the goal of methods based solely on exon counts is to find *differential* changes in relative exon expression across groups. This is based on the assumption that the sequencing-related biases would be the same across samples, which may not be true.

Methods that analyze relative exon counts to detect differential alternative splicing, like DEXSeq (Anders *et al.*, 2012) and the `diffSplice` method of voom (Law *et al.*, 2014), fit a linear model per gene to the counts per exon, allowing for an individual exon effect which quantifies the relative exon expression i.e. how different an exon is from the overall mean gene expression. Then they find differentially spliced exons by detecting exons who have different exon effects in the two groups. In fitting this model, standard count distributions like the negative binomial model, along with the corresponding shrinkage of the dispersion, can be used in the same manner as for gene expression. For this reason relative exon usage depends on global behavior of the gene, unlike the inclusion-exclusion approach which is quite local.

Like the inclusion-exclusion approach, relative exon usage also focuses on individual exons rather than global isoform estimates. However, the linear model also implies that the usage of the exon is defined relative to all other exons in the gene, which means that identification of an exon effect in the linear model does not directly translate to alternative splicing differences in that exon (Anders *et al.* (2012)). For example, if many of the exons in a gene are alternatively spliced then, relative to the mean, the unusual exons with large exon effects are actually the few that are *not* alternatively spliced. Similarly, introns should not be included in a relative exon usage analysis since they will overwhelm the gene model. Neither of these situations are a problem with the inclusion-exclusion framework, where the measurement of splicing given by the PSI is truly local and independent of other, non-adjacent, exons in the gene.

On the other hand, relative exon counts can find differential usage of exons that cannot have inclusion-exclusion data; for example exons at the beginning or end of a gene do not produce skipping reads even if some isoforms do not use those exons.

We note that in addition to targeting the correct exon, the paradigm of inclusion-exclusion offers one possibly significant advantage compared to an analysis of the relative usage of exon counts, regardless of the specific statistical method. In the inclusion-exclusion paradigm, those exons that show no reads skipping the exon in *any* sample of any group are naturally excluded by getting a p-value of 1 by definition (and similarly for those that show that fragments always skip the exon, such as introns). To get a sense of the value of using information about how many junction reads skip an exon, we can look at our real data example of 30 samples from 2 tissue types (described in Section 5.4). For each exon, our annotation of the isoforms that contain the exon results in a classification of an exon as constitutive (i.e. should not be excluded in any of the transcripts if the annotation is completely accurate) or as alternative (the exon is excluded in at least one isoform).

Of the constitutive exons (12.7% of the exons in the data), only 1% (529 exons) show *any* junction reads skipping the exon in *any* of the 30 tissue samples, while 35.0% of those exons annotated as alternatively spliced have such junction reads. This strongly suggests that the implicit removal of exons with no skipping junctions is preferentially removing exons that are not alternatively spliced, which ultimately can increase the power (Bourgon *et al.*, 2010). There is no natural way to exclude such exons in the linear model analysis of exon counts, since the constitutive exons are actually important in building the gene model – though a post-analysis filtering could be implemented to eliminate exons that were never skipped.

Clearly this implicit filtering can also be a disadvantage if many alternatively spliced exons are excluded because of a lack of sufficient reads to detect the skipping event. We view this as less of a practical disadvantage because we find that in practice practitioners are likely to want evidence in the form of junction reads skipping an exon in order to have faith in the call of an exon as alternatively spliced or to design an experiment to test the result. But a more general concern is that because the inclusion-exclusion paradigm relies heavily on reads that span the junctions of exons, which are a small percentage of all reads, it relies on a lower number of reads and could have lower power.

## 3 **Modeling the Dispersion**

There are several different distributional choices for a dispersion model for the binomial. A very common choice is the beta-binomial model which places a beta prior on the standard binomial distribution proportion parameter. This distribution is not a member of the exponential family and there is not a closed-form solution for the MLE. This model has been used recently in the setting of methylation data (Sun *et al.*, 2014; Feng *et al.*, 2014; Dolzhenko and Smith, 2014), but less so in that of gene expression settings (with Zhou *et al.* (2011) and Hardcastle and Kelly (2013) being exceptions). In a related fashion, the MATS method of Shen *et al.* (2012) creates a dispersed model by placing a uniform prior on the proportion parameter of the binomial which is a specific example of a beta prior where the beta parameters (and thus dispersion) are set in advance; unlike the other methods previously cited, MATS was actually developed for the setting of measuring PSI and detecting alternative splicing from mRNA-Seq data.

Another common approach is to use quasi-likelihood methods; the estimates for the proportion (mean) remain the same as that found from a binomial model, but the distribution of the estimate of the mean depends on a dispersion parameter and thus results in greater estimates of variability in the presence of over-dispersion (or less if under-dispersed) than the standard binomial model in the case of a quasi-binomial. An existing method of analyzing proportions in a genomic setting in this fashion is the modified extra-binomial (EB2) method of Yang *et al.* (2012) which follows an alternative quasi-likelihood approach given in Williams (1982) where the variance is aligned to match that of a beta-binomial; again this was not developed for differential exon splicing, but for comparing allele frequencies between populations.

We focus our methods on the double exponential family (Efron, 1986), which is a full probability model that results in estimates closely related to the quasi-likelihood method. This class of distributions, which we will describe in detail below, adds a dispersion parameter to any member of the exponential family. This distribution has the advantage of being closely related to the quasi-likelihood approach and yet still provides a likelihood platform for the development of shrinkage methods. Furthermore, the distribution is itself in the two-parameter exponential family, making calculations and approximations straightforward. Because our method of shrinkage for dispersion estimates applies beyond just the binomial distribution, we will describe the development in general terms using the entire double exponential family of distributions rather than concentrating on the binomial setting (which we will refer to as the “double binomial” distribution).

**Notation** In what follows, the data from every exon consists of an entry *Y*_*ij*_ and *m*_*ij*_. *Y*_*ij*_ refers to the count of the number of times there was inclusion of the event *j* for sample *i*. For the setting of exon inclusion, *Y*_*ij*_ would be the number of fragments overlapping exon *j*. *m*_*ij*_ gives the total possible number of counts related to event *j*; in the exon setting this would be the total number of fragments given by the sum of those expressing exon *j* and those skipping exon *j*. The value *Y*_*ij*_/*m*_*ij*_ is the “percent spliced in” (PSI) value and is the standard binomial estimate of the probability of inclusion. For concreteness, we will assume that the PSI is per exon, with the understanding that the same methods could be applied to other ways of defining “included” and “excluded” fragments. The *m*_*ij*_ we will call the total count, meaning the total number of reads both including and skipping the exon.

For the rest of this section, we will focus on the modeling of the distribution of *Y*_*ij*_ for just a single exon *j*, and therefore we will drop the subscript *j* when the meaning is clear.

### 3.1 **The double exponential family of distributions**

The double exponential family of distributions of Efron (1986) generalizes any distribution in the exponential family of distributions by including an over-dispersion parameter. Specifically, assume that the naive choice for the distribution of *Y*_*i*_ is a distribution in the exponential family, such as Binomial or Poisson. We also assume that for each *Y*_*i*_ there is a corresponding *m*_*i*_, which might be all equal in some settings. For the binomial, *m*_*i*_ is clearly the total number of trials. For other distributions, *m*_*i*_ might be taken as 1 for all samples. We follow the notation of Efron (1986) so that we transform *Y*_*i*_ into a random variable *Z*_*i*_ whose mean *μ* under the naive distribution is independent of the sample size *m*_*i*_, e.g. *Z*_*i*_ = *Y*_*i*_/*m*_*i*_ for the binomial family. We write the pdf of the naive distribution in canonical exponential family form by

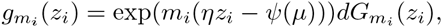

where *η* = *η*(*μ*) is the link function relating the mean *μ* to the canonical parameter *η*, and *ψ*(*μ*) is the normalizing function. For the case of binomial, the link function is the standard logit function, *η*(*μ*) = 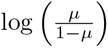 and the normalizing function *ψ* is given by *ψ*(*μ*) = *-* log(1 *− μ*). We can also define the standard deviance residual of *Z*_*i*_ from the mean *μ* in terms of the canonical parameters of the exponential family,

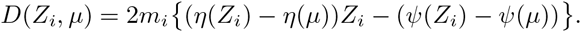

Note that *D*(*Z*_*i*_, μ) is the deviance based on the naive (non-dispersed) model.

Efron (1986) proposes adding a dispersion parameter *ϕ* to any distribution in the exponential family so that the the dispersed distribution has a pdf given by

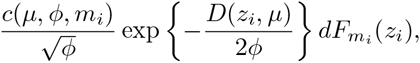

where *c*(*μ, ϕ, m_i_*) is a normalizing constant. The role of *ϕ* is reminiscent of the role of the variance parameter in a normal distribution where *ϕ >* 1 implies over-dispersion and *ϕ <* 1 implies under-dispersion.

**Approximating** *c*(*μ, ϕ, m_i_*) The normalizing constant *c*(*μ, ϕ, m_i_*) can be computationally expensive to calculate, especially in the context of genomic studies where the maximization routines will need to be calculated for every exon. Efron shows that

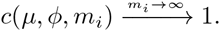

Note that this approximation is as *m*_*i*_ is large. In our exon setting *m*_*i*_ is the total number of reads for the exon in a particular sample and exon – including both the reads overlapping the exon and the junctions skipping the exon – and is *not* the number of samples.

Approximating *c*(*μ, ϕ, m_i_*) with 1 results in the mean of *Z*_*i*_ remaining approximately *μ* while the variance of *Z*_*i*_ from the over-dispersed distribution becomes approximately

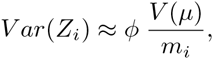

where *V* (*μ*) is the variance function for the naive (non-dispersed) distribution from the exponential family. If the naive distribution is the binomial distribution, the corresponding over-dispersed distribution has variance approximately 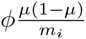, while for a Poisson, the over-dispersed distribution has variance approximately *ϕμ*.

**Comparison of Double Binomial to Beta-binomial** By analogy, the beta-binomial distribution can be parameterized in terms of the mean *μ* and dispersion parameter *ρ ∈* (0, 1), which represents the correlation between the *m*_*i*_ Bernoulli draws. Then the variance of *Z*_*i*_ from a beta-binomial is given by

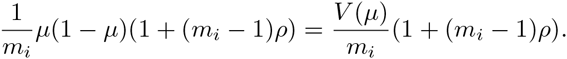

Note that beta-binomial only allows for over-dispersion, unlike the double binomial, though that is not of great concern in most genomic studies where over-dispersion is expected.

We compare their density functions in Supplemental Figure S1 after aligning their first two moments (i.e. choosing *ρ* so that *ϕ* = (1 + (*m*_*i*_- 1)*ρ*)). Not surprisingly, for low levels of dispersion, the double binomial and beta-binomial are quite similar since they both converge to a binomial distribution as the dispersion vanishes. For large dispersion parameters or for the mean near the boundary of values of 0 and 1, they show the greatest differences. When the dispersion is extremely large (log(*ϕ*) = 3), the double binomial puts more mass on the boundary values compared to the beta-binomial. This is a rare dispersion value in the data we are examining (Supplemental Figure S2), though when we look at the number of zeros in real data, it matches simulations from a double-binomial more closely than that of a beta-binomial (Section 4.3). More applicable for our data is the differences between the two when *μ* is near 0 (or 1), where the beta binomial concentrates the mass much closer to 0 while the double binomial allows for more variability.

Over the rest of the range of parameters, their distributions are relatively similar, but the distribution of the beta-binomial is more difficult to work with analytically than the double binomial, particularly when we use the approximation of *c*(*μ, ϕ, m_i_*) = 1 for the double binomial.

**Multiple Groups** For applications, we are interested in the case where there are covariates that results in different *μ* for different samples *i*. We focus on the common case in genomic studies where the covariates simply define *K* separate groups of samples and we want to compare the mean proportion between the groups for every exon. The most common example of this is the two group comparison where *K* = 2. For example the two groups might be disease and normal samples, or samples with a gene knocked-out versus those without; more generally any setting where there is comparison of treatment samples versus control samples. Higher values of *K* can be found, for example, in time course data, where each group might correspond to observations observed at the same time point, or just generally where there are multiple sets of treatments for each sample.

In the case of multiple groups, we have a separate mean *μ_k_* for each group *k*; when the meaning is clear we can refer to this vector of means as *μ*. Maximizing the joint likelihood with the approximation that *c*(*μ_k_, ϕ, m_i_*) = 1 gives simple analytical MLE estimates for *μ* and *ϕ*,

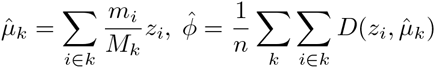

where *M*_*k*_ = ∑*i ∈ k m_i_* is the total coverage of all samples in group *k*. Thus the approximation *c*(*μ_k_, ϕ, m_i_*) = 1 results in the standard estimates of *μ_k_* from the non-dispersed distribution and estimates of the dispersion based on deviance residuals. These estimates are the same as those in the quasi-likelihood setting, giving a likelihood-based method that closely resembles the results of the quasi-likelihood method (Efron, 1986).

This approximation leads to enormous computational savings as well as alignment with the standard quasi-binomial approach, and the methods we develop rely on this approximation.

### 3.2 **Distribution of 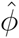**

A key component of our methods of shrinkage is that we can find a statistic that relies solely on the parameter *ϕ* and is independent of *μ*. This allows us the opportunity to shrink the estimates of *ϕ* based on shared information across exons independently of the value of *μ* for a particular exon.

Distributions from the double exponential family are also members of the two-parameter exponential family. This implies that the conditional distribution of 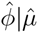 defines a conditional likelihood independent of *μ* and that the conditional distribution can be easily approximated using a modified profile likelihood (see Pawitan (2001) for a review). This conditional likelihood can be used for shrinkage of the estimates 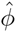 as in Robinson and Smyth (2007).

However, if we further make use of the approximation *c*(*μ_k_, ϕ, m*) = 1, it can be shown that 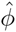 is asymptotically independent of the vector of sample means, 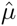 Furthermore, 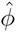 is asymptotically Gamma distributed,

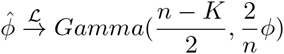

where 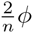 is the scale parameter of the Gamma (see Supplementary Text, Section 1).

This result does not depend on the form of the naive distribution that formed the starting point of the analysis, though we will use the binomial distribution since we are analyzing percentages (PSI). For example, it could similarly be used for an over-dispersed Poisson distribution for gene expression analysis, which we discuss in the conclusion. However, in this setting, unlike the more commonly used negative binomial distribution, the relationship between the mean and variance would be linear, not quadratic.

## 4 **Development of Empirical Bayes Methods**

We now will develop the methodology underlying our two shrinkage methods for the dispersion, as well as test statistics for comparing differential skipping between groups.

Our methods rely on an empirical bayes strategy for shrinkage of the dispersion parameter. Empirical bayes strategies are widely used, for example in, EBSeq (Leng *et al.*, 2013) and DSS (Wu *et al.*, 2013) for differential gene expression detection. In addition, empirical bayes methods are used in a wide range of problems, in particular for gene expression analysis of microarrays where the widely used limma method (Smyth, 2005) provides empirical bayes shrinkage of the variance parameter of a linear model. We develop an empirical bayes method for the family of double exponential distributions, which we call DEB-Seq (Double exponential Empirical Bayes with application to Sequencing).

We also consider the weighted likelihood approach which is implemented for the negative binomial distribution in the widely-used gene expression method `edgeR` (Robinson and Smyth, 2007). We show that when applied to the approximate likelihood of 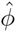 based on the double exponential family of distributions, it gives an estimator of a similar form as our empirical bayes estimator, except with a different parameterization of the prior distribution. To distinguish the method with this form of parameterization we refer to this as the WEB-Seq method (Weighted Empirical Bayes shrinkage with application to Sequencing).

We also compare our methods to other existing methods that provide shrinkage of the dispersion parameter in comparing proportions across groups. These methods perform shrinkage of dispersion in slightly different ways. The modified extra-binomial shrinkage method (EB2) (Yang *et al.*, 2012) was developed to test for differences in allele frequencies, though it can be easily applied to the setting of differential exon usage. The method employs shrinkage by reparameterizing the variance function in terms of two global parameters that are estimated via linear regression by combining data across all SNPs. MATS (Shen *et al.*, 2012), mentioned above, was developed for detecting differences in PSI and assumes a uniform prior for the proportion parameter of a binomial and further adds a correlation between the two conditions being compared; this parameter is assumed shared by all the exons and thus provides implicit shrinkage. BBSeq (Zhou *et al.*, 2011) is a method developed for gene expression studies that shrinks the dispersion parameter of a beta-binomial model. Their shrinkage method fits a cubic polynomial to the independently estimated, logit-transformed dispersion parameters as a function of the fitted values of the observed data. The method of Feng *et al.* (2014) provided in the `DSS` package was developed for DNA methylation data and also provides shrinkage of the dispersion parameter of a beta-binomial distribution. They do so by fitting an empirical bayes model that assumes the dispersion parameter has a log-normal prior distribution and their computations are based on method of moments estimators for the beta-binomial. For convenience, we will refer to this as the DSS method, though DSS actually refers to the corresponding gene expression technique that the same authors developed earlier in Wu *et al.* (2013).

### 4.1 **Double exponential Empirical Bayes**

Empirical bayes estimation of the dispersion parameters via an explicit likelihood formulation is a natural way to provide shrinkage estimators of the dispersion parameter. By this we mean the following general strategy for estimating a parameter *θ*: formulate a Bayesian model *Y |θ ∼ F* and *θ ∼ G_α_*, let the estimate 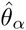 be given by *E*_*α*_(*θ|Y*), and then choose the parameter *α* by estimating *α* from the marginal distribution of *Y α*. This results in an estimate 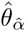 which has the desirable shrinkage properties of Bayesian estimators but also has an explicit method for estimating the parameter *α*.

Many distributions, including the distributions in the double exponential family, do not have a prior that gives a tractable form for the marginal distribution of *Y* to permit easy estimation of *α* from the data. However, if we make the approximation that the normalizing constant *c*(*μ_k_, ϕ, m_i_*) is equal to 1, we show in Section 1 of the Supplementary Text that 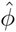 is approximately independent of the group means and Gamma distributed with known shape parameter (*n − K*)*/*2, and scale parameter equal to 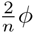

This suggests a simple bayesian model of *ϕ_j_* for each exon *j*. A conjugate prior for the scale parameter of a gamma distribution with a known shape parameter is the inverse gamma distribution, *IG*(*α*_0_, *β*_0_) ^1^. Applying this conjugate prior to the scale parameter *ϕ_j_*, we can formulate the Bayesian model

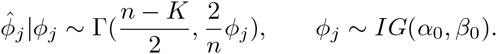

This gives the posterior distribution of 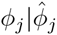. as a Inverse-Gamma distribution with shape parameter 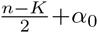 and rate parameter 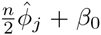. Then a standard Bayesian point estimate of *ϕ_j_* is given by the mean of the posterior distribution,

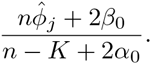

To give an empirical bayes solution, we estimate *α*_0_ and *β*_0_ from the marginal distribution of the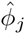. across all exons, making the assumption that the/inline/are independent across exons. Critically, the approximate distribution of 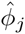. is independent of the individual total counts, *m*_*ij*_, per exon and sample. This means that under the Bayesian model above, the 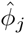. are marginally identically distributed and we can use the joint likelihood of 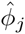. to find estimates 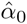. and 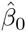. Because the Inverse Gamma prior for *ϕ* is a conjugate prior this results in an analytical expression for the marginal density of 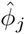. given by the generalized beta distribution (Raiffa and Schlaifer, 1961) with density,

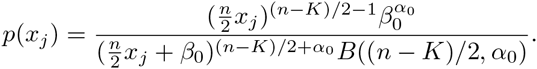

We estimate 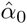. and 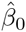. by maximum likelihood estimation.

Then the empirical bayes estimate of *ϕ_j_* is given by

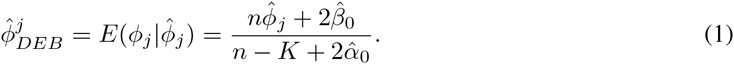

We call this estimation procedure Double exponential Empirical Bayes with application to Sequencing (DEB-Seq).

### 4.2 **Empirical Bayes based on Weighted likelihood**

Wang (2006) gives a general strategy for combining likelihoods by creating a (weighted) average of the log-likelihoods of all the experiments. Robinson and Smyth (2007) adapt this idea to the gene-expression setting to make it gene-centric, and this procedure is implemented in the `edgeR` package. Each gene *j* is given a separate weighted log-likelihood 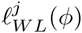 which is the weighted sum of the individual gene log-likelihoods for gene *j*, 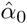 and a common log-likelihood which is the sum of the individual gene log-likelihoods for all genes *£_CM_* (*ϕ*),

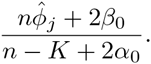

Then for each gene *j*, the shrunken estimate 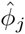 is given by maximizing 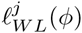. The weight *δ* given to the common log-likelihood *£_CM_* (*ϕ*) is a tuning parameter that must be chosen. McCarthy *et al.* (2012) suggest that it be chosen so that it is proportional to sample size adjusted by degrees of freedom, with the `edgeR` package assigning a fixed value for *δ* in the gene expression setting by default equal to 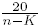

In order to apply this to the dispersion parameter of a negative binomial, Robinson and Smyth (2007) use the conditional likelihood of 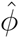 given the sufficient statistic 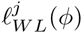 to define the log-likelihood *£^j^*(*ϕ*). For the negative binomial, this requires that the total count parameter *m*_*i*_ be equal for all samples, which is not the case in typical RNA-Seq experiments, so Robinson and Smyth (2008) provide a method of getting pseudo-totals by implementing what they call a quantile-adjusted conditional maximum likelihood procedure (qCML). Essentially, the observed data is adjusted via an iterative algorithm to simulate pseudo-data that is distributed from a negative binomial with equal library sizes.

We can follow the same weighted likelihood strategy to create a weighted likelihood for a distribution from the double exponential family. Again, if we assume that *c*(*μ_k_, ϕ, m*) = 1 we have that 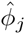is approximately distributed 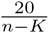. Unlike the negative binomial distribution, the approximate likelihood of *ϕ_j_* does not depend on the *m*_*ij*_, eliminating the need to create pseudo-data in the implementation. Furthermore, the resulting estimate from maximizing the weighted log-likelihood has an analytical solution unlike the case of the negative binomial,

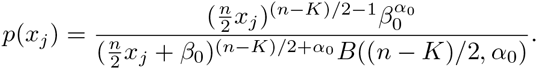

Where 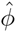. is the average of the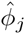. over all exons in the data.

**WEB-Seq** One major advantage of the empirical bayes method in estimating the dispersion is that the amount of shrinkage performed is entirely determined from the data, unlike the weighted likelihood method. It is not known, especially in the context of exon usage, if a single default *δ* will perform adequately across a range of differing experiments.

It is clear from comparing the weighted likelihood estimate with that of the empirical bayes estimate that the weighted likelihood estimate takes on the same form as that of the empirical bayes estimate in our case where we use dispersed distributions from the double exponential family. Specifically,

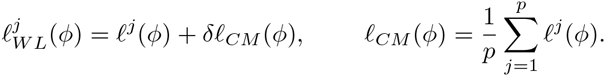

This implies the weighted likelihood method can be written as an empirical bayes solution where the prior is parameterized by a single variable *δ* rather than the two parameters *α*_0_ and *β*_0_, (We are implicitly treating 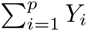. as a fixed value, rather than explicitly conditioning on it, but 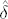. will normally be the average of thousands, if not tens-of-thousands, of exons). Note that in the weighted likelihood approach, *δ* is assumed to be strictly positive, and 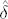. will similarly be positive, therefore satisfying the assumptions for *α*_0_ and *β*_0_ to yield a true density.

This naturally suggests an estimator based on this alternative parameterization to fuse these two methods together, which we call a Weighted Empirical Bayes shrinkage with application to Sequencing, or WEB-Seq. Using this parameterization and maximizing the density as described above for the empirical bayes method gives us an estimate 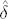 and represents the amount of shrinkage that should be performed in the weighted likelihood method as determined by the data. The resulting dispersion estimates are then,

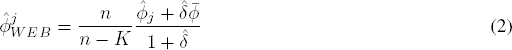

(for details concerning estimation of *δ* see Supplementary Text, Section 2).

We will see that WEB-Seq has similar performance to the original empirical bayes approach, though it is more conservative and as a result slightly less powerful. Both methods perform well, and we choose to focus on this method largely because it appears to be more robust in simulations to violations of the model due to being more conservative (Section 5).

### 4.3 **Estimates of under-dispersion near the boundary**

We found in initial simulations that estimated proportions lying on the boundary (i.e. either exactly 1 or 0, corresponding to an exon that is either never skipped or always skipped in the sample) have a large and adverse effect on the false discovery rate (FDR) as the sample size increases (see Supplementary Figure S3). This is particularly detrimental for the exon skipping application, since many exons may be close to being never skipped or never expressed, particularly if introns are included in the analysis.

This behavior is because the method estimates under-dispersion for such exons, leading to a large number of false discoveries. Ironically, the effect is worse with larger sample sizes: the effect becomes noticeable around 5-10 samples per group and for increased sample sizes the FDR grows without control. The reason for the increase with larger sample sizes is that exons with proportions all 1 or all 0 across *all* samples get a p-value of one, which results in an implicit filter of the data. For exons whose true proportion is near the boundary, the observed data is more likely in low sample sizes to have observed proportions entirely on the boundary and therefore assigned a p-value of one and removed from consideration. In larger sample sizes, there is an increased chance that a non-boundary sample will be observed, allowing the exon to remain in the analysis and have an effect on the FDR results.

There are several ad hoc approaches to this issue. One is to filter exons with mean proportion close to the boundary. While a successful filtering procedure can result in a positive impact on the power of a test (Bourgon *et al.*, 2010), in this case it is unsatisfactory to have a test-statistic that is so sensitive to the degree of filtering. Another approach is to not allow under-dispersion by setting the dispersion in such cases to one, i.e. binomial variance (see also the most recent version of DESeq that does not allow the dispersion estimate to be decreased via shrinkage).

We obtained better results by defining an effective sample size, *n*_*eff*_, defined as the maximum of the number of non-boundary samples and *K* + 1, where *K* is the number of groups being compared. We use this *n*_*eff*_ to adjust the degrees of freedom, *n −K*, that appears in the Gamma prior, changing it to be *n*_*eff*_ − *K* instead. Note that this means that each exon has a slightly different effective sample size. A similar difference in effective sample size can result when a sample has *m*_*ij*_ = 0 in which case the sample cannot be included in the analysis of that exon (see Supplementary Section 2 for details).

The result of using these effective degrees of freedom is that it becomes less likely to erroneously estimate under-dispersion on the boundary, though it remains possible to estimate under-dispersion. With this adjustment no under-dispersion is in fact estimated at any sample size for any exon in our simulated or real data sets, while previously exons with proportion parameters near the boundary were frequently estimated to be under-dispersed and thus given inflated significance. Further, a continuous range of dispersion values *ϕ* are estimated for these boundary exons which we find to be more natural than forcing them all to have no dispersion, which would be the case if we arbitrarily set those with under-dispersion estimates to have 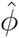 = 1.

In Supplementary Table S1 we compare the proportion of exons that are affected by our adjustment to the degrees of freedom, for data simulated under either a double binomial or beta-binomial distribution, as well as the real data. We see that 78% of the exons are affected by these changes, with 43% having a large reduction of 5 or more. This highlights the importance of boundary control in the exon skipping problem. Interestingly, we see that the real data more closely follows the double binomial simulations in this respect, rather than the beta-binomial.

### 4.4 **Comparison of conditions**

After obtaining estimates of *ϕ* for each exon based on shrinkage across the exons, we then return to our question of testing differences of inclusion between conditions. These can be formulated in the form of contrasts on the vector *η*, and we can test the significance of the contrast per exon. In our implementation, we focus on the common two group comparison, though all of the methods carry through to the more general setting of contrasts of groups. In the setting of comparing two groups, we reparameterize so that *β*_1_ = *η*_2_ − *η*_1_ is the value of the contrast defined by the difference of the two groups (and *β*_0_ = *η*_1_). We test the null hypothesis *H*_0_ : *β*_1_ = 0.

Let 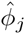^*^ be an estimator of *ϕ* (e.g. DEB-Seq or WEB-Seq, or the MLE 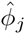). Then for testing the specific null hypothesis *β*_1_ = 0, the standard GLM approach defines the likelihood ratio statistic as

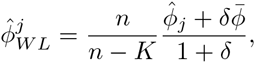

where *S* and *S*_*H*_0 are one half of the sum of the deviance residuals of the full or null model, respectively (i.e. 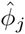 for 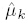 estimated under the full or null model, respectively) and 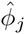^*^ is the estimate of *ϕ* under the full model. For WEB-Seq or DEB-Seq, we use the shrunken estimates of *ϕ* given by the shrinkage methods outlined above 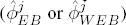.

Because our shrinkage methods give analytical solutions for the estimates of *ϕ*, we can easily compare the effect of using the shrinkage estimator 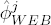 instead of the original MLE 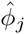 on the statistic *W*. 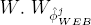 will be smaller than 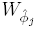 if 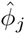 is less than 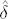 the mean of the 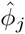 across all exons. Therefore, those exons with small estimates of variability will become less significant after shrinkage. If the distribution of 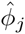 *is* roughly gamma this implies that the majority of test statistics are reduced in significance since the gamma distribution is skewed right and therefore the median is less than the mean.

Asymptotically, *W*_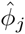_ should follow a *F* distribution with (*K-* 1) and (*n −K*) degrees of freedom (Jorgensen, 1997). Based on our simulations, we find that this approximation is poor for small sample sizes, e.g. when the size of each group is five or less, leading to poor control of Type I errors. Instead we propose an alternative statistic, which re-estimates *ϕ* under the null and alternative, that has much better performance in small sample sizes,

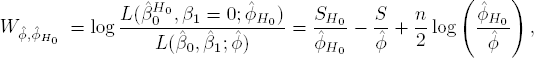

This means the likelihoods are not strictly nested, so that 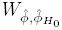 can technically take on negative values. Here, 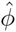_*H*_0__ and 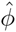 are estimates of *ϕ* based on the null (*K* = 1) and the alternative (*K* = 2) models, respectively. For our shrinkage estimates of *ϕ* in equations (1) and (2), this means that the value of *K* and the unshrunken estimate 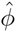 change. We also re-estimate the shrinkage values of 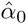, 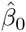 and 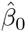 which are based on all exons’ unshrunken estimates of *ϕ* estimated under the assumption that *K* = 1 or *K* = 2 accordingly.

We also find that the shrinkage methods result in less variability in the estimate of 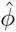 with the result that the likelihood ratio statistic more closely follows a 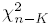 distribution than the standard *F* distribution for unshrunken estimates.

## 5 **Evaluation of Methods**

We evaluate WEB-Seq and DEB-Seq on simulated and real data sets, and compare them to other methods currently existing in the literature.

### 5.1 **Description of the Simulation**

Exon count simulations were created under a two group comparison setting. For the purpose of imitating real data, we chose simulation parameters based on fitting models to 170 Acute Myeloid Leukemia samples generated by the Cancer Genome Atlas project (Cancer Genome Atlas Research Network, 2011), see Supplementary Text, Section 3 for details. We generated data under a double binomial distribution and also a beta-binomial distribution for evaluation of the robustness of our techniques, all of which were developed assuming the data come from a double binomial distribution. Either 1% or 10% of the exons were randomly selected to show differential usage between the groups; for these non-null exons, the treatment effect, *β*_1_, was picked uniformly from the union of [-3,-0.5] and [0.5,3], accounting for both decreased and increased exon usage. Otherwise, exons were given a treatment effect of 0, comprising our null set of exons. For each simulation, 85,373 exons were simulated and a basic filtering process was applied to remove exons with proportions all equal to 1 or all equal to 0 across the samples (i.e these result in p-values of 1).

We used the simulated data to evaluate the methods developed above: 1) the empirical bayes method with a single parameter prior (WEB-Seq) 2) the general two-parameter empirical bayes method for the prior parameters (DEB-Seq), and 3) the weighted likelihood method with *δ* fixed to be equal to the default value implemented in `edgeR` 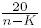. In addition to our dispersion-shrinkage methods, we implemented the shrinkage methods of BBSeq, EB2 and DSS, briefly described in Section 4.1. The MATS method described earlier does not take as input inclusion and exclusion count matrices, but rather creates its own from BAM alignment files, and thus could not be compared on the simulated data.

We also implemented three methods that fit a dispersion parameter per exon but with no shrinkage across exons: quasi-binomial GLM estimation as implemented in the `glm` function in R (R Core Team, 2013), maximum likelihood estimation based on a beta-binomial distribution implemented using the `betabin` function from the `aod` package in R, and maximum likelihood estimation based on an approximate double binomial distribution where the normalizing constant is set to 1. The quasi-binomial GLM and the double binomial MLE are closely connected, as described in Section 3.1, and are both non-shrinkage counterparts to our methods. However, the quasi-binomial estimation by default uses Pearson residuals to estimate the dispersion, rather than deviance residuals. The beta-binomial maximum likelihood method is the non-shrinkage counterpart of the BBSeq and DSS methods that rely on the beta-binomial distribution.

For each method, the estimation routines were performed and the p-values were adjusted to control the FDR to a 0.05 level using the standard Benjamini-Hochberg FDR procedure (Benjamini and Hochberg, 1995) implemented in the `p.adjust` function in R (R Core Team, 2013). The final measures of performance were the methods’ ability to control false discoveries and their power to detect non-null exons over the 100 simulations.

### 5.2 **Comparison of double binomial methods on simulated data**

We first compare the performance of WEB-Seq and DEB-Seq to other methods that also rely on the double binomial distribution, particularly those without shrinkage. As hoped, the shrinkage methods improve upon the estimation of the dispersion parameter, reducing the mean squared error (MSE) significantly from the unshrunken versions of the double binomial (Supplementary Figure S5). DEB-Seq had the smallest MSE, followed by WEB-Seq, and both were a significant reduction compared to double binomial estimation methods with no shrinkage.

We compare their ability to control the false discovery rate across a range of sample sizes for a fixed FDR cutoff (Figure 2) and see that WEB-Seq and DEB-Seq were also superior in control of FDR across a range of samples sizes. For data simulated as double binomial, they both control the FDR well, while unshrunken methods have high rates of FDR compared to the target 5%. Weighted likelihood shrinkage for the double binomial with a pre-determined tuning parameter (based on `edgeR` default) is erratic in its control of FDR for small sample sizes (5 or less in each group), but then adequately controls FDR. However, the pre-determined tuning method becomes over-conservative for large sample sizes and the result is a large drop in power for weighted likelihood for large sample sizes.

**Figure 2:**
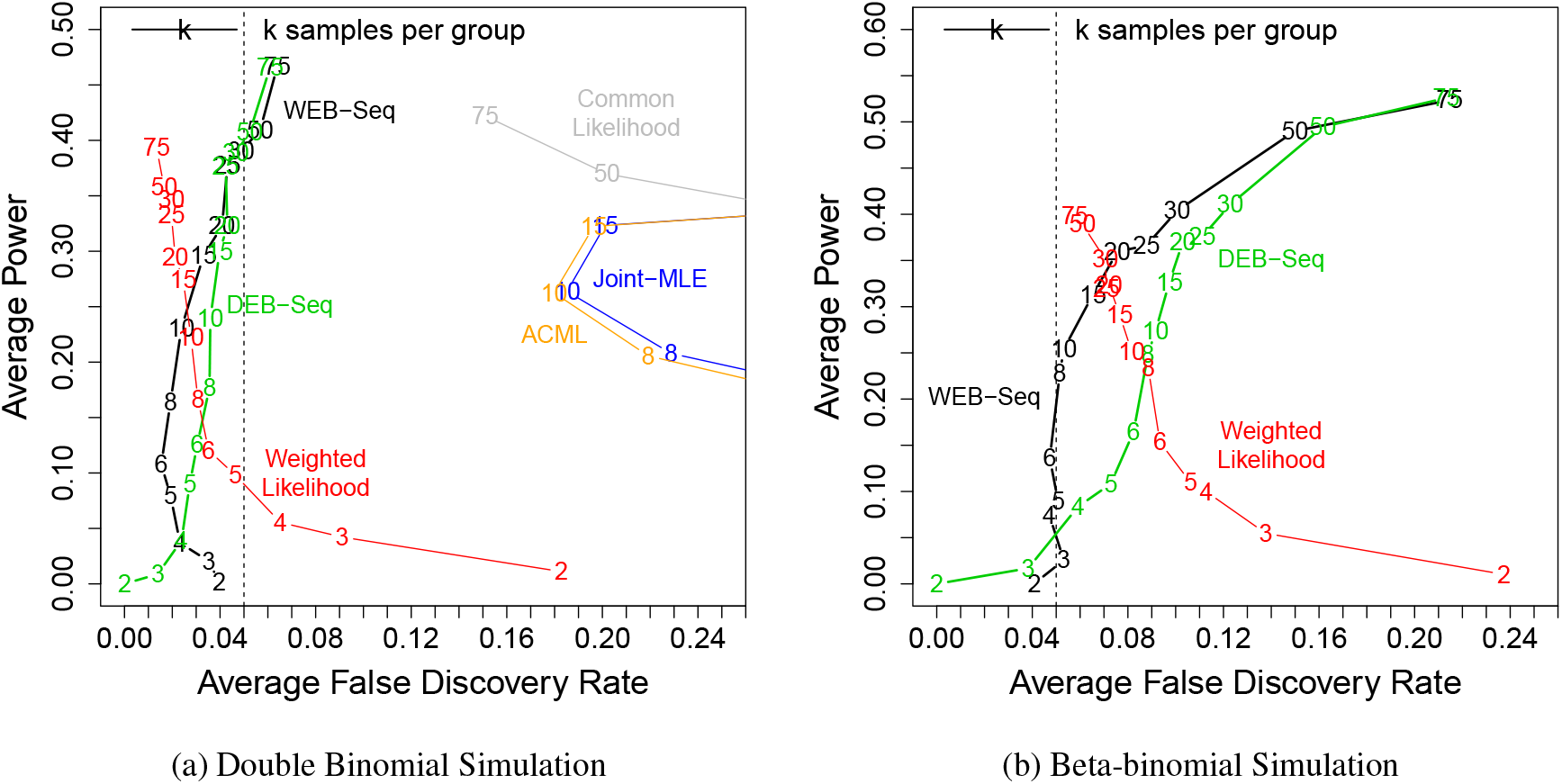
**Double Binomial Methods compared across different sample sizes for a fixed 5% FDR cutoff**. Plotted is the average Power (y-axis) against FDR (x-axis) over various sample sizes across 100 simulated data sets with 1% of exons differentially used, based on p-values adjusted to provide a 5% FDR level. The results for a single method across different sample sizes are connected by a line. The numbers that overlay a method denote the power and FDR for that specific sample size (*per group*) in a 2 group comparison. The 5% FDR boundary is given by the dotted vertical line. The data are simulated under (a) a double binomial distribution and (b) a beta-binomial distribution. The methods shown are all based on a double binomial to account for over-dispersion: 1-parameter empirical bayes (WEB-Seq); 2-parameter empirical bayes (DEB-Seq); weighted likelihood shrinkage with fixed parameter; estimation of a single dispersion parameter *ϕ* for all exons (common likelihood); the double binomial ML estimate (Joint-MLE); and the approximate conditional ML estimate (ACML). The joint MLE, ACML and common likelihood are not shown in the beta-binomial simulation because their FDR values were beyond the limits of the plot.

To evaluate the robustness of the methods, we consider data that do not follow the double binomial distribution, but rather the beta-binomial distribution. Again, in small sample sizes (4 per group or less) both WEB-Seq and DEB-Seq control the FDR accurately. For moderate sample sizes (10 per group or less), the more conservative WEB-Seq maintains control of the FDR; DEB-Seq still has greater power, but has an increase of FDR to about 10% for these moderate sample sizes. For large sample sizes (10 or more per group), there also appears to be an underlying bias due to the p-values being calculated under the wrong model so that the FDR of both WEB-Seq and DEB-Seq start growing well beyond the 5% target.

### 5.3 **Comparison to other methods on simulated data**

**FDR Control** In Figure 3, we compare how WEB-Seq controls the FDR in small sample sizes (5 samples per group) compared to other existing methods. WEB-Seq shows superb control of the FDR, while all of the other methods – both those that use shrinkage and those that do not – have FDR rates far beyond what their adjusted p-values would indicate.

**Figure 3:**
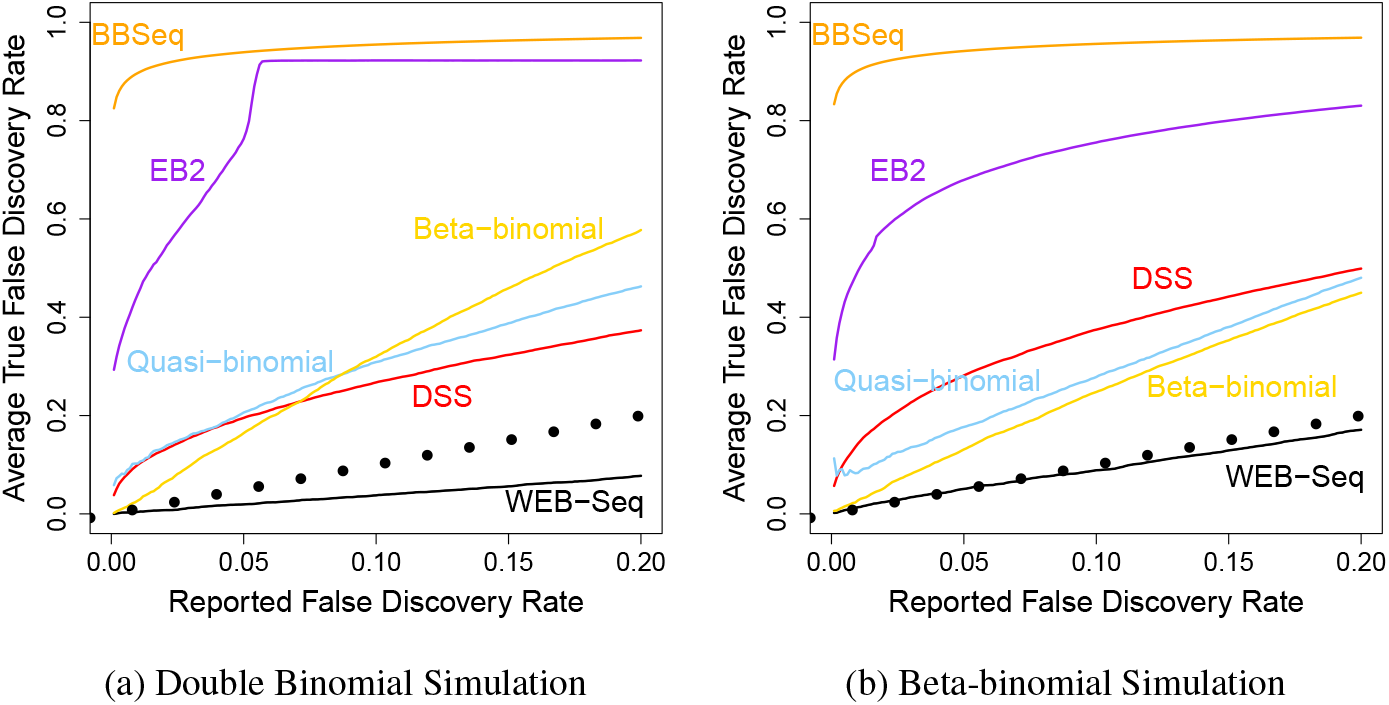
**Control of FDR:** Plotted is the true FDR level against the reported FDR for different methods, indicated by different colors. The reported FDR refers to the cutoff used for calling exons significant based on FDR-adjusted p-values. The true FDR is calculated for those exons called significant by knowing which exons are truly false discoveries in our simulations. The dotted line is the *y* = *x* line. The data are simulated under (a) a double binomial distribution and (b) a beta-binomial distribution. Here the percent of non-null exons was 1%. For 10% non-null and other sample sizes, see Supplementary Figures S7-S14.

This pattern holds for different distributions of the data (double binomial or beta-binomial), for different choices of sample sizes, for unbalanced sample sizes, and for different choices of the percent of non-null exons (Supplementary Figures S7-S12). All of the specific numbers given in what follows are assuming a 1% non-null exons; the same lack of control hold for 10%, though with different specific values.

Among the alternative methods, the beta-binomial MLE with no shrinkage performs the best in controlling the FDR under both count distributions, but still has an FDR much larger in small sample sizes than that indicated by their adjusted p-values, even when the data is distributed according to a beta-binomial distribution (e.g. 10-15% FDR instead of the target rate of 5%). The performance is even worse for double binomial distributed data, indicating a lack of robustness to the modeling assumptions.

Quasi-binomial (also with no shrinkage) appears to perform similarly to the beta-binomial MLE in Figure 3, but evaluations of other sample sizes (Supplementary Figure S6-S10) shows that its FDR performance is erratic and can be much larger than its target value; similarly, the double binomial GLM (not plotted) fails to control the FDR at any sample size we explored and converges to an FDR at around 40%.

The remaining methods that perform shrinkage (EB2, BBSeq, and DSS) also rely on beta-binomial dispersion models, but do quite badly in FDR control even for beta-binomial data. The true FDR rates for BBSeq and EB2 range from 0.96 and 0.98 (*n* = 3) to 0.27 and 0.41 (*n* = 75) when the target FDR control was 0.05. DSS is slightly better since it eventually controls the FDR, but it only does so starting at sample sizes that are quite large for genomic studies (*>* 20 per group); for *n* = 3 and *n* = 5 per group, the true FDR rates are 0.53 and 0.28 respectively, when the target rate is 0.05.

In contrast, WEB-Seq controls the FDR at the desired level for the double binomial data and beta-binomial data in small sample sizes, and only shows poor control of the FDR for fairly large sample sizes (*>* 10 per group), and even then only for beta-binomial distributed data.

Due to multiple testing corrections, the comparative performance of the various methods hinges on the distributional behavior of their test statistics far in the tails of the distribution. At standard levels of individual (per exon) Type I error control (e.g. 0.01-0.05), the p-values of all of the methods do a reasonable job of controlling the false positive rate. However, after controlling for multiple testing, the effective raw p-value cutoff for significance at a target FDR rate of 5% is around 0.0001 if only 1% of the exons are non-null (and 0.001 if 10% are non-null). In this tail of the distribution, the test statistics can perform quite differently with respect to Type I error control and their resulting power. In our simulations, WEB-Seq controls Type I error even far into the tails of the distribution, unlike any other method (Supplemental Figures S25-S28) indicating that it is the performance of the methods in the tail of the distribution (and not the p-value adjustment method) that leads to this discrepancy.

We would also note that for exon analysis, the number of exons evaluated can be quite high compared to gene expression analysis (easily 50,000-100,000 exons), and in many studies we would not expect the percent of exons that are skipped emphdifferentially between groups to be very large: even 1% translates to hundreds of differentially spliced exons. With an even smaller percentages of true non-nulls, which could be common in real studies, control of the FDR will become even worse for these other methods.

**Power** For comparison of the power of the methods, we must similarly focus on the small levels of Type I error relevant for multiple testing corrections. Traditional ROC curves show that WEB-Seq results in greater power than all the other methods except DSS when the data comes from a double binomial distribution; for data distributed under a beta-binomial, it still has slightly greater or equivalent power to all the methods except DSS (Supplementary Figures S15 − S18). This again illustrates the robustness of WEB-Seq to the modeling assumptions. DSS is the only method that has clearly improved power, mainly in small samples sizes (e.g. power of 3% for WEB-Seq versus 8% for DSS for 3 samples per group); however, as we have seen DSS also shows very poor control of the FDR for those sample sizes.

Many biological studies focus on the top performing exons for validation and followup analysis, especially when there are large numbers of significant results; this is clearly related to power, but specifically oriented to top ranking exons. We find that the WEB-Seq method also provides p-values that do well at prioritizing the truly non-null exons in small sample sizes. However, the DSS method is also competitive in raw rankings, which is not surprising given the general power performance discussed above. In Figure 4, we plot the average proportion of false discoveries in the top-ranked exons for simulations with *n* = 3 samples per group. We see that for this small sample size, the alternative methods have a much higher proportion of false positives in the top-ranked exons compared to WEB-Seq, regardless of whether the data is distributed beta-binomial or double binomial. For slightly larger sample sizes (e.g. 5 vs 5), however, DSS outperforms our method in ranking the exons. Beta-binomial MLE performs clearly worse in rankings for small sample sizes, even when the data is distributed according to a beta-binomial. For much larger sample sizes (e.g. 10 vs 10), the difference between the DSS, beta-binomial, and WEB-Seq methods is minimal and they all perform better than any of the other competing methods (Supplementary Figures S19-S22).

**Figure 4:**
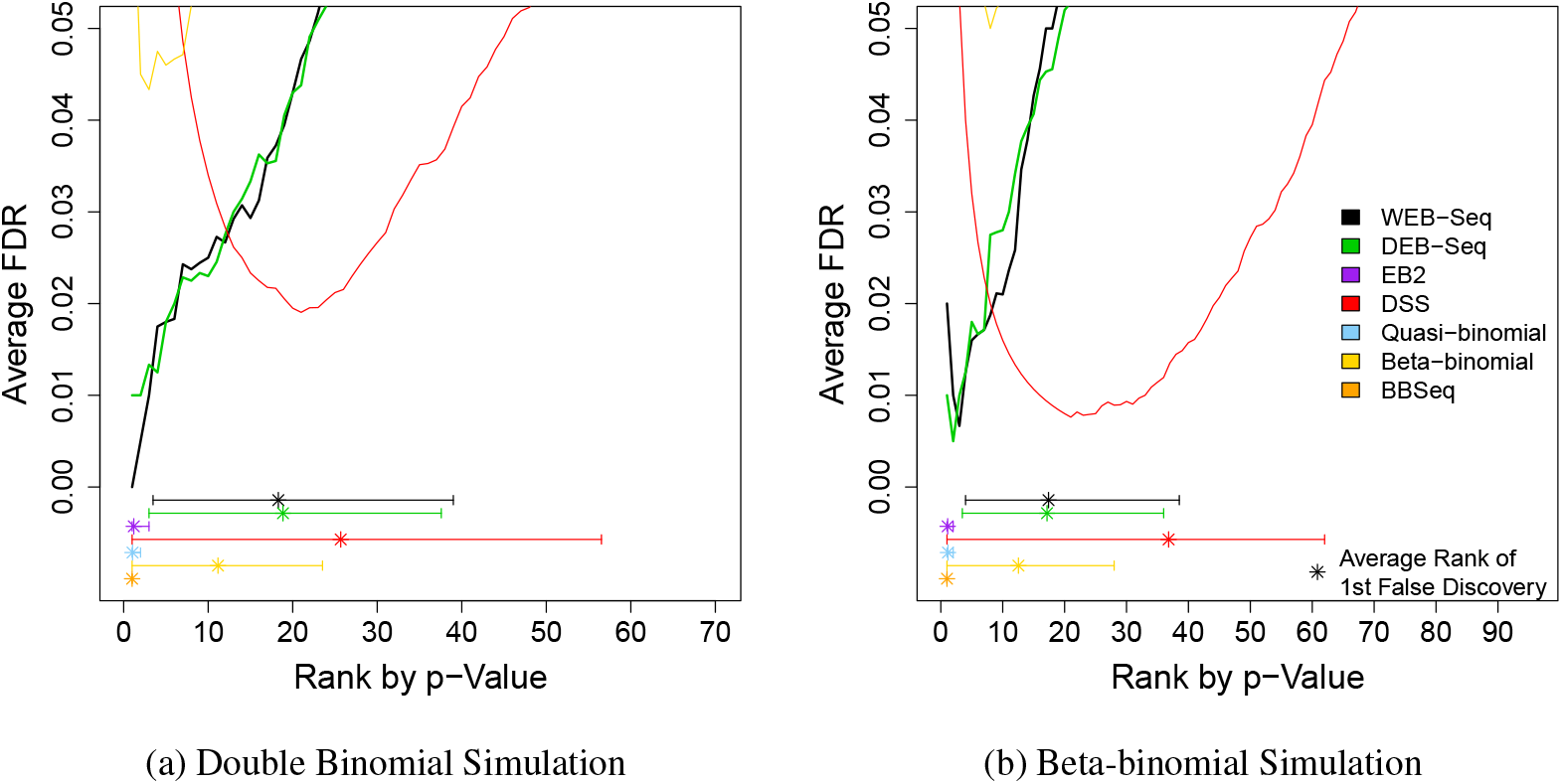
**False Discoveries by Rank.** Plotted is the average proportion of false discoveries (y-axis) in the top *x* exons (x-axis) for a 3 versus 3 comparison with 1% of exons alternatively spliced. For each method, the 2.5% and 97.5% percentiles of the rank of the first false discovery across the 100 simulations is given by the horizontal lines, with the average rank of the first false discovery marked by an asterisk.

In summary, WEB-Seq is much superior in giving accurate FDR adjusted p-values across the ranges of sample sizes that correspond to those frequently seen in practice, while every other method performs poorly, and often dramatically so. WEB-Seq also shows high power compared to the other methods and performs well at prioritizing exons. For some ranges of sample sizes the DSS method has more power than WEB-Seq, but at those same sample sizes, DSS has highly inaccurate p-values for assessing significance. The main competitor in small sample sizes when considering both FDR control and power is the beta-binomial MLE with no shrinkage, but it still has poor FDR control, less power, and worse rankings of the exons in small sample sizes – even when the data is actually simulated from the beta-binomial distribution.

### 5.4 **Comparison of methods on TCGA data**

In order to have a reasonable setting for detecting differential alternative splicing we downloaded RNA-Seq data from two different tumor types also sequenced by the TCGA: Stomach and Ovarian. For comparisons between these two sets of tumors, we expect that there should be large differences in alternative splicing due to the simple fact that the tumors originated from two different tissue types, and tissue-specific alternative splicing is well documented (Pan *et al.*, 2008). We note that the differences in tissue types is a rather extreme example since in this case we do expect a large number of significant exons. The RNA-Seq data was realigned using TopHat v1.4.1 (Trapnell *et al.*, 2010), and exon inclusion and exclusion counts were calculated for exons annotated by ENSEMBL version 66 (Flicek *et al.*, 2013). See Supplementary Text, Section 4 for more details about the processing of the data.

We create a ‘null’ situation to compare the methods, where the two groups that are compared are both of the same tissue type. We note that these are tumor samples, so there may be differential alternative splicing in the different tumors even though they are the same tissue type, but since these samples are randomly assigned to the two groups this is unlikely to be a significant factor.

We ran the double binomial methods, as well as the other methods on the TCGA data sets (Table 1). MATS could only be run in the null setting, as the stomach and ovarian samples were of different read types and the MATS software did not support this.

We compare the proportions of exons called significant in the null setting across methods based on FDR adjusted p-values. Note that this is not a measure of FDR nor traditional Type I error; in fact, if we believe that there are no significant exons, *any* discoveries in an all-null setting would technically imply that the rate of false discoveries is 1.

Focusing on just our double binomial based shrinkage methods, WEB-Seq and DEB-Seq, we see that these call essentially 0% of exons significant in the null case (see Supplementary Table S2). Examining individual simulations rather than the average of 100 simulations shows that indeed there are truly zero exons called significant the majority of the time in every sample size. As in the simulations, WEB-Seq has slightly less ‘power’ than DEB-Seq in that it makes fewer significant calls under the real setting where we compare the different tissue types.

Turning to the alternative methods, the average false positive rates (based on FDR corrected p-values) from the null analysis in Table 1 suggest BBSeq and EB2’s poor control of the FDR in the simulated data is echoed in the real data. Across the samples sizes, EB2 finds roughly 7% of the exons significant and BBSeq finds 4-6% significant in the null setting. DSS shows better performance in the null analysis with less than 1% false calls (except for *n* = 2); but in the exon setting where there are many exons to evaluate, this still ranges from around 700 exons (3 vs 3) to 170 exons (7 vs 7) incorrectly called significant compared to only 4 and 1 exon calls for WEB-Seq in those same sample sizes, respectively (see Supplementary Tables S4 and S3). For comparison with MATS, we applied WEB-Seq to the inclusion-exclusion count matrices produced by MATS. In the null setting, MATS appears to have a call rate between 1.8% and 3.9% (646 to 1,557 exons called significant), while WEB-Seq makes at most one call for any given sample size.

These false positive rates based on FDR adjusted p-values do not directly give FDR rates like those in the simulations, and because it is a null setting where there are no truly significant values, the FPR should technically be exactly 0 to achieve FDR control. For comparison, if 10% of the exons were truly non-null and the method had 100% power, the percentage of false positives would have to be 0.6% to get an FDR at the target level 5%; 1% of exons non-null would require the percentage of false positives to be. 05%. And in practice the false positive rate would need to be *much* lower since no method has anywhere near 100% power (in fact the power is quite low for these small sample sizes). A 3-7% false positive rate would then mean a minimum FDR of 21-38% and in practice much higher. This indicates that the lack of control of FDR shown in our simulations appears to be supported by implementation on the real data.

**Table 1:** **Comparison to Alternative Methods.** Shown in the table below are the average percentage of exons called significant based on FDR corrected p-values from the Tissue Data under 100 simulations of the null and real scenarios described above. DEXSeq was post-filtered to have the same set of exons as the inclusion-exclusion setting. For all the results shown below, except for MATS, the total number of starting exons is 412, 002 but the rates are percentages out of only those exons that had at least one skipping event, a number which varies with sample size but is roughly 1/4 of all exons. The results from MATS are based on a different set of exon data produced internally by MATS, roughly 35,000 exons; WEB-Seq results are not shown on this set of exons, but WEB-Seq makes at most one significant call on the MATS set of exons (for sample sizes 3, 5 & 7) and zero for other sample sizes. Furthermore, MATS was run on a single random sample because of the time involved in processing a single run of the data (see Section 5.6). See Supplemental Table S4 for the precise number of exons called and the results from the non-shrinkage methods.

| Sample Size | DEXSeq |  | EB2 |  | BBSeq |  | MATS | DSS |  | Beta-bin. |  | WEB-Seq |  |
| --- | --- | --- | --- | --- | --- | --- | --- | --- | --- | --- | --- | --- | --- |
|  | Real | Null | Real | Null | Real | Null | Null | Real | Null | Real | Null | Real | Null |
| 2 vs 2 | 13.50 | 1.46 | 11.08 | 7.47 | 18.78 | 6.56 | 3.45 | 10.43 | 1.31 | 5.06 | 0.00 | 0.89 | 0.00 |
| 3 vs 3 | 17.17 | 0.62 | 11.56 | 7.20 | 24.95 | 7.37 | 1.77 | 12.19 | 0.58 | 9.70 | 0.00 | 2.82 | 0.00 |
| 4 vs 4 | 21.77 | 0.12 | 11.94 | 6.85 | 28.90 | 6.20 | 2.45 | 14.50 | 0.19 | 13.73 | 0.00 | 6.73 | 0.00 |
| 5 vs 5 | 26.63 | 0.08 | 12.67 | 6.54 | 31.28 | 6.02 | 2.74 | 17.37 | 0.16 | 17.73 | 0.00 | 12.16 | 0.00 |
| 6 vs 6 | 30.00 | 0.11 | 13.27 | 6.26 | 32.62 | 5.73 | 3.93 | 19.44 | 0.18 | 20.41 | 0.01 | 15.94 | 0.00 |
| 7 vs 7 | 33.86 | 0.06 | 14.08 | 6.00 | 33.82 | 4.98 | 3.39 | 22.03 | 0.14 | 23.42 | 0.00 | 20.26 | 0.00 |

The only other method that gives comparable results to WEB-Seq is the beta-binomial MLE method. However, since the needed false positive rate based on adjusted p-values has to be so small in many real settings, it is difficult to compare these two on just these evaluations. Instead, we directly compare the false positive rate based on the uncorrected p-values in the null setting as we vary the cutoff (see Figure 5 and Supplementary Figure S29); we see that the performance directly echoes that seen in the simulations (Supplementary Figures S25-S28). Only WEB-Seq (and DEB-Seq) control Type I error in the tails of their distributions for small sample sizes. As in the simulations, beta-binomial is the next best, but still has an increase in its Type I error comparable to that of the simulations, which resulted in FDR rates of around 15% rather than the target 5%. All of these analyses strongly suggest that the problems we see in controlling the FDR in simulations are real and will appear in analyzing real data sets.

**Figure 5:**
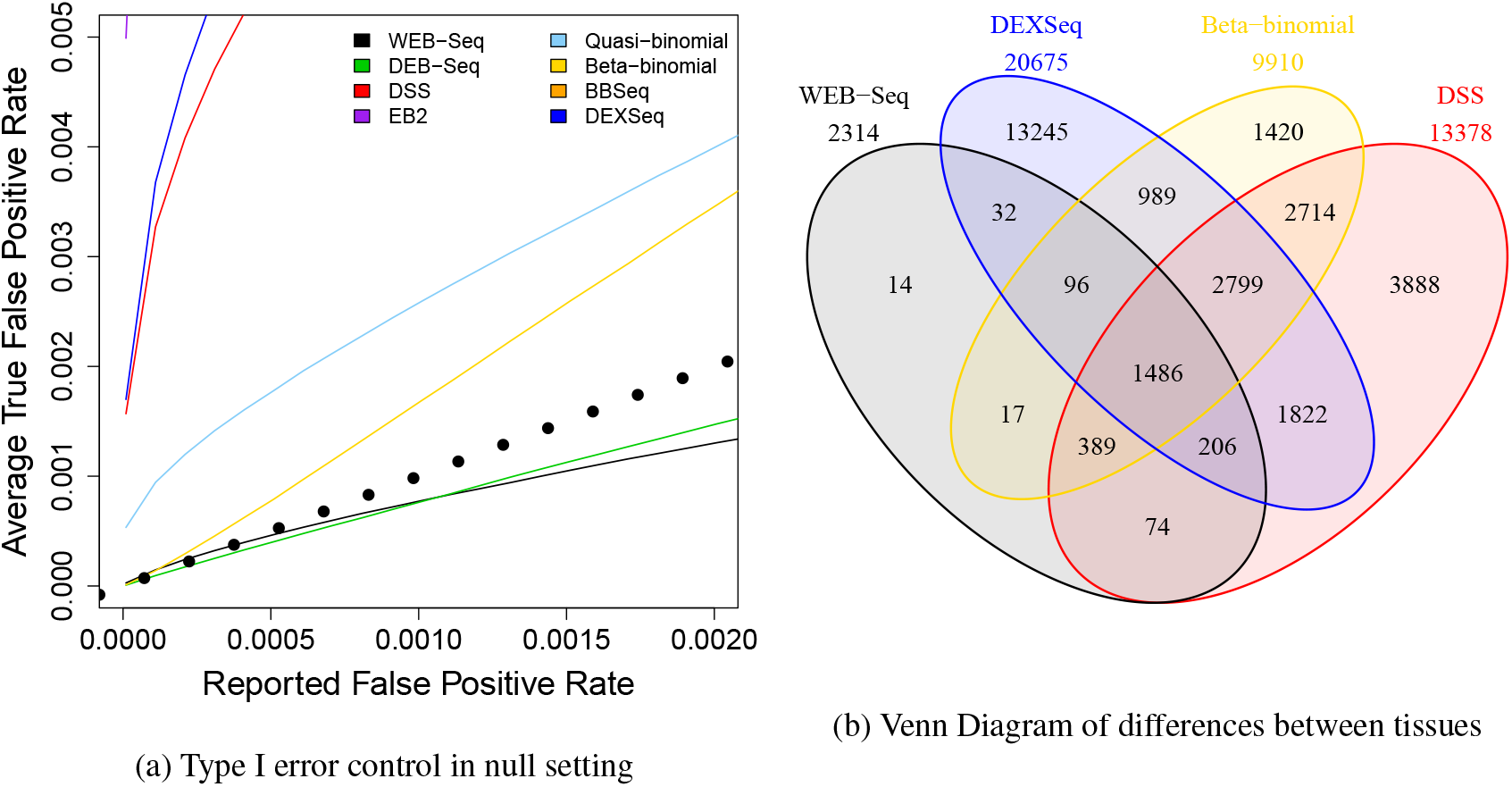
**Evaluation on TCGA data,** *n* = 3. (a) Type I error on of TCGA “null” setting where we plot the percent of exons found significant in the null setting as a function of the p-value cutoff based on uncorrected pvalues. (b) We show Venn diagrams of the analysis of differences between tissues for WEB-Seq (black), DEXSeq (blue), Beta-binomial MLE (yellow), and DSS (red). DEXSeq is limited to only those exons also found to have skipping events and so also considered by the other methods. Numbers in the Venn diagrams are based on overlapping counts averaged over the 100 simulations.

For the comparison in the real setting where the two different tissue types are compared, we see many more calls made by the other methods compared to WEB-Seq. Given the problems these methods have in obtaining accurate FDR control in both the null setting and our simulations, we expect that some portion of these additional calls in the real setting are due to a much higher level of false discoveries than reported. We compare the overlap in these calls via Venn diagrams (Figure 5 and Supplementary Figure S29), and we see that almost all of the calls made by WEB-Seq are also found significant by another method. If we look at the overlap of WEB-Seq with the six alternative methods we considered here (DEXSeq, DSS, EB2, BBSeq, beta-binomial MLE, and quasi-binomial), we see that 85% of WEB-Seq’s calls are supported by at least *four* other methods for a 3 samples per group comparison (Figure 6 and Supplementary Figures S30-S31). Beta-binomial MLE is the next best candidate yet only has 20% of its calls so strongly supported, in spite of the fact that two of the other methods (DSS & BBSeq) are also based on the beta-binomial distribution.

**Figure 6:**
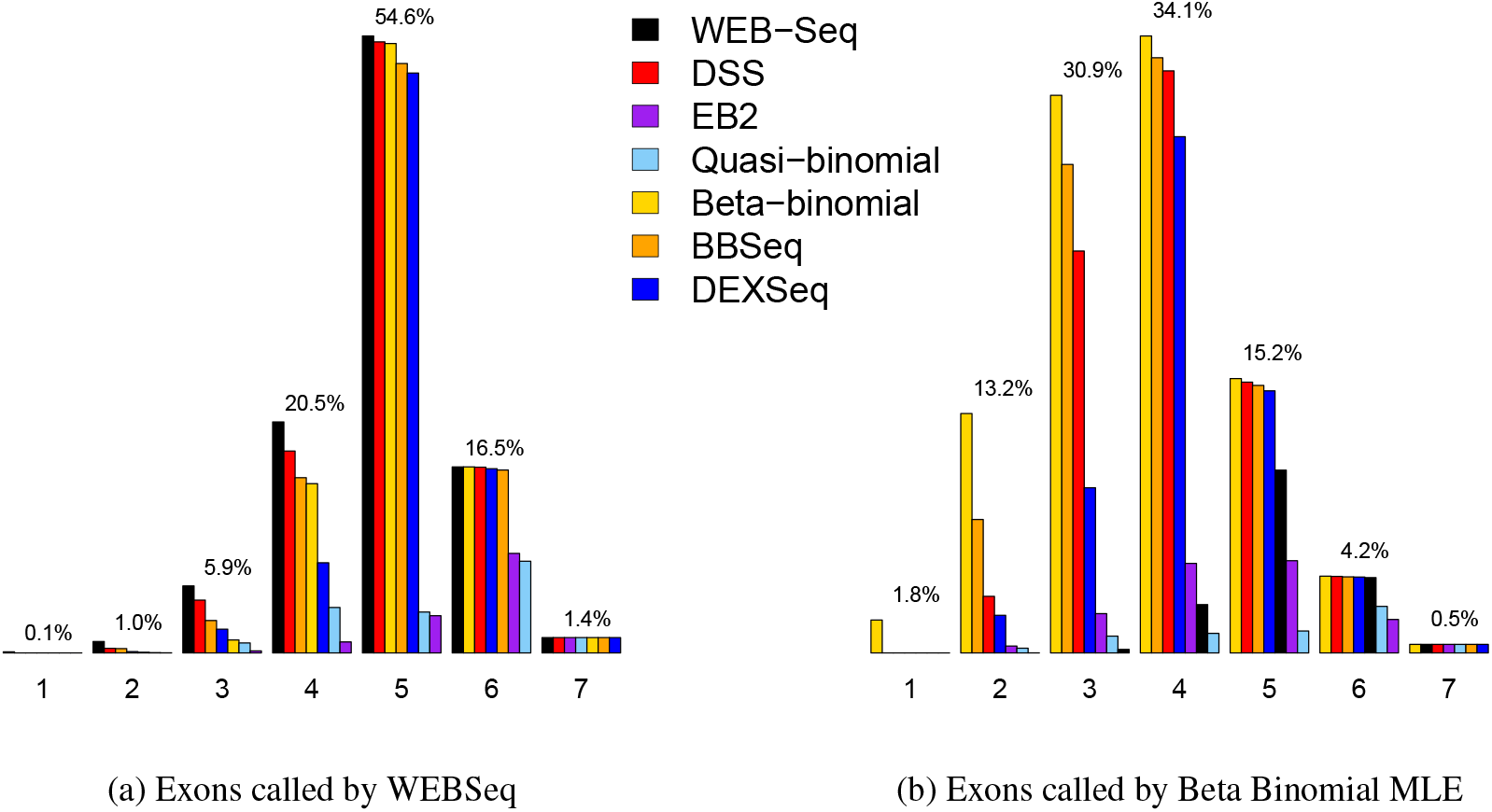
**Percentage overlap with calls by other methods,** *n* = 3. (a) For WEB-Seq we calculate what percent of significant calls made by WEB-Seq were called significant with the other six methods. An exon called significant by WEB-Seq could be called significant by one to seven other methods, one signifying only WEB-Seq called it significant, while seven indicates that all methods call the exons significant. Plotted are the percentages called significant for 1-7 other methods (given as both black text as well as a barplot corresponding to the method’s color). Within each category of number of overlapping methods, we further indicate which of the other methods are actually overlapping, with a barplot indicating the percentage of those exons that are called significant by each method, colored according to corresponding method. Only for overlap=2 will the percentages of the other methods add up to the overall overlap percentage. The figures for the seven methods are split over two pages. In (b) we repeat the exercise for exons called significant by beta-binomial MLE. For all seven methods and for *n* = 6, see Supplementary Figures S30-S31.

### 5.5 **Comparison to relative exon usage**

We made a further comparison of the performance of our method to another popular method of finding differential alternative splicing in exons, DEXSeq (Anders *et al.*, 2012). The DEXSeq method relies on the relative exon usage framework described above in Section 2.3 and *only* uses exon counts, without using information about how the junctions skip exons. Therefore, DEXSeq is not just an alternative statistical method, but also a fundamentally different summarization of the mRNA-Seq data as compared to the inclusion-exclusion setting for which our methods are developed. For these reasons, it is not clear that you can make a reasonable comparison between WEB-Seq and DEXSeq. However, DEXSeq is a popular method for detecting alternative splicing with just exon counts, so we attempt some basic comparisons.

As we discussed in Section 2.3, relative exon usage evaluates all the exons, not just those showing skipping. Because of the difference in exons evaluated between the methods, we concentrate on comparing the performance of DEXSeq for just those exons that show skipping in the samples being analyzed; this means we run DEXSeq on all 412,002 expressed exons, as required by the algorithm, and subsequently filter down to the set of exons that show junction reads, and then finally calculate adjusted p-values based only on these exons. For this set of exons the false positive rate of DEXSeq is similar to that of DSS, particularly for the smaller sample sizes (Table 1), as is its control of the Type I error rate on the ‘null’ setting (Figure 5 and Supplementary Figure S29). This suggests that DEXSeq will have similar problems as DSS in having much higher rates of false discoveries than indicated by the adjusted p-values.

DEXSeq clearly calls more exons significant in the real setting than any of the methods based on junction counts, even when limited to the same set of exons. This could be a sign of increased power when using the exon counts, as mentioned above. When we look at the Venn diagrams in Figure 5, we see that this increase constitutes tens of thousands of additional exons called significant, and 30%-40% of the calls of DEXSeq are not supported by any of the junction count methods, regardless of sample size (Supplementary Figures S30 − S31). Furthermore, as we explained in Section 2.3 (and has been noted by the authors of DEXSeq), relative exon usage does not always reliably indicate the right exon within the gene that has differential usage. The additional calls may also be a reflection of this problem, as well as increased power.

We also consider DEXSeq’s performance more broadly and consider their calls on all exons, even those without skipping reads. About 12% of constitutive exons are called significant by DEXSeq (Table S6). This is roughly their total representation in the data so DEXSeq does not appear to be preferentially finding exons annotated to be alternatively spliced. If we instead compare only exons that are the sole exon called significant in their gene, following the theory they are more likely to be targeting the correct exon, even a larger percentage are annotated as constitutive *and* furthermore have no reads skipping them in the data for *any* of the 30 samples (18%-46%, Table S6). In comparison, in WEB-Seq, 0.2% of the significant exons (or 76 exons) are annotated as constitutive and all of them, by definition, have reads skipping them to at least justify the call of significance.

We can also evaluate the data properties of the significant exons to evaluate whether they demonstrate data characteristics that would lead us to trust the call. We compare the density of the log-Fold-Change between the groups of the odds-ratio of skipping an exon for the significant calls made by both methods (Supplementary Figure S24). WEB-Seq clearly has a much stronger tendency to find exons with large differences in the skipping proportion, which is not surprising given that that is the basis of its test statistic, unlike DEXSeq. More striking is that for DEXSeq there are significant peaks at 0, indicating many of the exons found significant by DEXSeq do not show evidence of differential exon usage in the form of a difference in the proportion of skipping counts. The constitutive exons found by DEXSeq, in particular, are completely centered at zero. This could be because of the lack of identification of the correct exon, explained above; when we examine the “single-exon” genes which are presumed to target the appropriate exon, these exons show slightly greater propensity to be removed from zero (Supplementary Figure S23c).

Ultimately, we find the inclusion-exclusion paradigm concentrates the analysis on those exons with tangible evidence of alternative splicing as well as directly highlighting the specific exons of interest. We suspect this will also be an effective way of preventing a large source of false discoveries as well as being robust to the behavior of the other exons in the gene.

### 5.6 **Computation Time**

In an exon usage analysis, the method used needs to potentially be able to handle all exons, a number which for the human genome can be in the hundreds of thousands. Under the inclusion-exclusion paradigm there is an automatic filter of exons with either no skipping reads or no overlapping reads, which significantly reduces the set of exons under analysis. After this automatic filter we see ranges between 40K and 200K exons in this scenario for real RNA-Seq experiments when looking at just protein coding exons. Because we have exact analytical solutions for our estimators of WEB-Seq, the only numerical optimization involves the calculation of the estimates of the prior parameters of the gamma distribution. In a 5 versus 5 setting, analyzing a total of 105,275 exons, WEB-Seq required only 25 seconds on a single core computer running an AMD Opteron 6272, 2.1 GHz processor. WEB-Seq can be run for any size experiment on a single core, personal laptop in under a minute.

Some of the other methods that we compared require much greater computation time. BBSeq is time intensive requiring 4.3 hours to complete the analysis; calculating the MLE for beta-binomial for each exon similarly took 2.8 hours. Quasi-binomial took 30 minutes, and EB2 and DSS only require 2 minutes. With the MATS algorithm, it is difficult to compare computational times directly since it is only implemented in a format that requires processing of all of the raw sequences to create the counts, which in the 5 vs 5 example took a total of 35 hours, as well as needing approximately 150GB of available space for intermediate files.

For the relative exon usage example of DEXSeq, filters of the exons are not appropriate and therefore the number of exons being analyzed range between 300K and 400K exons for the human genome, while the inclusion-exclusion setting automatically filters out those without any skipping reads and results in roughly 100K exons. Analyzing the full set of the 412,002 expressed exons took the newest version of DEXSeq 4.5 hours to complete on a single core (DEXSeq allows the use of multiple cores, but for comparison purposes we restricted it to a single core). When we artificially set the number of exons to be the same 105,275 analyzed in WEB-Seq, DEXSeq required 1.1 hours compared to 25 seconds using WEB-Seq, but we note this was for timing comparison only, since filtering them in this way would invalidate the DEXSeq analysis.

## 6 **Discussion**

We have developed a novel method for providing shrinkage estimators for the dispersion parameter of a dispersed exponential family of distributions. We rely on a dispersion model that is closely connected to the common quasi-likelihood method for providing over-dispersion to a binomial, which are widely used and numerically robust. By making use of the distributional form of Efron (1986), we have shown that there is a simple formulation of the approximate distribution of the dispersion parameter and that this form provides a straightforward empirical bayes method to estimate shrinkage. In effect, we provide a likelihood-based empirical bayes method for quasi-likelihood estimation of the dispersion parameter. By further relating this empirical bayes method to weighted likelihood shrinkage methods (Robinson and Smyth, 2007), we give a non-standard parameterization of the Gamma prior that leads to an alternative estimator in this class of estimators that demonstrates some areas of improved performance. Further, our distributional form and the empirical bayes method that results do not require any tuning parameters, unlike the weighted likelihood methods of `edgeR`.

In comparison to other methods for analyzing exon-skipping events, we showed in both simulated and real data that our method is the only one that can accurately control the FDR in the sample sizes that are commonly seen in genomic studies (often less than 5 samples per treatment group), with the other methods having very large false discovery rates compared to their reported rate. We also show that our method has good power and the ability to prioritize truly significant genes.

For detecting differential alternative splicing, we have discussed that there are alternative summaries of the data that require different statistical techniques than those presented here. The percent spliced in statistic may not be the most appropriate for every setting. We compared directly to the relative expression approach – using only exon counts without using the information in the junction reads – and we illustrated that reliance on junction reads naturally filters the problem to those exons most likely to be differentially used. Another alternative approach relies on estimating the expression levels of individual isoforms, and this may give more insight into alternative splicing particularly when there is a great deal of information about the transcriptome that is being sequenced. However, in our experience there are still many cases where researchers find themselves without a well constructed annotation of the transcriptome and would have to rely on de-novo methods to construct genes and/or transcripts. This is an extremely complicated problem, and these de-novo methods can be unreliable and unstable if used on a single, small experiment or without significant depth (see also Anders *et al.* (2012) for a discussion of isoform versus exon analysis). In contrast, inclusion-exclusion counts rely on detection of exons and splice sites, which are much simpler problems. In short, inclusion-exclusion counts provide useful, interpretable information about the undergoing of alternative splicing within the organism, and our method gives a reliable technique for the statistical analysis of such data. While our shrinkage method is quite general, we have focused on our motiving example, detecting differential usage of exons between conditions in order to detect group-specific alternative splicing. In particular, our data examples were drawn from mRNA-Seq data, and the simulations were based on parameters estimated from that same data. There are other settings that require the comparison of a large number of proportions between groups, for example in the setting of comparing allele frequencies or differential methylation, and it is possible that the performance would differ in those settings due to differences in the properties of the data. Similarly, our evaluation of the shrinkage method is based on the binomial distribution as our starting point because of our interest in exon inclusion rates, but our entire methodological development is completely general and can be applied to any distribution from the exponential family. While every type of data should have careful development for its unique properties, it is useful to have a single framework that can be the starting point for so many settings. Possible examples could be that of analyzing differential gene expression data (based on the Poisson distribution) or differential proportions of isoforms (based on multinomial distribution). Indeed, we tested a Poisson-based version of our WEB-Seq on gene expression data along with fourteen other gene expression techniques from the literature and found that on simulated data that our method performed well compared to the other methods across the range of distributions we tried. WEB-Seq ranked the significant genes better than most other methods (Supplemental Figures S32,S33). Its control of the FDR was also better than many common methods, particularly in small sample sizes (Supplemental Figure S34), but definitely not giving accurate FDR control like we presented in the exon setting (with true FDR for 5 vs 5 of around 7% rather than target of 5%). Furthermore, these conclusions hold true with data simulated under the negative binomial distribution which is the distribution for which most of the other methods (except ours) were developed. The only methods that did equivalent or better in *both* FDR control and power were `voom` (Law *et al.*, 2014) and `baySeq` (Hardcastle and Kelly, 2010); notably the popular `edgeR` (Robinson *et al.*, 2010) and DESeq (Anders and Huber, 2010) methods did not do as well in either power or control of FDR. We find this particularly encouraging for the general use of our shrinkage method in other settings, since we did not change or adjust the method in any way for the gene expression setting, other than switching the choice of distribution within the exponential family.

In summary, our method gives reliable and robust improvement to the analysis of exon splicing, which is a straightforward but important approach to analyzing the complicated structure of alternative splicing. Furthermore, because the shrinkage ideas apply generally to exponential family of distributions and have close links to the common GLM approach for analyzing data, it has the potential to be relevant for other applied problems.

1 recall that if *X* ∼ *IG*(*α*_0_, *β*_0_) this implies that 1/*X* ∼ Γ *IG*(*α*_0_ where *β*_0_ is the rate parameter

## Acknowledgements

We wish to thank Christopher Paciorek of the Berkeley Statistical Computing Facility for helpful input into the coding of the algorithms and parallelization of the comparisons. The results published here are in part based upon data generated by The Cancer Genome Atlas pilot project established by the NCI and NHGRI. Information about TCGA and the investigators and institutions who constitute the TCGA research network can be found at http://cancergenome.nih.gov/.

*Funding:* This research was partially funded by NIH grant U24 CA143799 and NSF grant DMS-1026441.

